# Newcastle disease virus (NDV) expressing the spike protein of SARS-CoV-2 as vaccine candidate

**DOI:** 10.1101/2020.07.26.221861

**Authors:** Weina Sun, Sarah R. Leist, Stephen McCroskery, Yonghong Liu, Stefan Slamanig, Justine Oliva, Fatima Amanat, Alexandra Schäfer, Kenneth H. Dinnon, Adolfo García-Sastre, Florian Krammer, Ralph S. Baric, Peter Palese

## Abstract

Due to the lack of protective immunity of humans towards the newly emerged SARS-CoV-2, this virus has caused a massive pandemic across the world resulting in hundreds of thousands of deaths. Thus, a vaccine is urgently needed to contain the spread of the virus. Here, we describe Newcastle disease virus (NDV) vector vaccines expressing the spike protein of SARS-CoV-2 in its wild type or a pre-fusion membrane anchored format. All described NDV vector vaccines grow to high titers in embryonated chicken eggs. In a proof of principle mouse study, we report that the NDV vector vaccines elicit high levels of antibodies that are neutralizing when the vaccine is given intramuscularly. Importantly, these COVID-19 vaccine candidates protect mice from a mouse-adapted SARS-CoV-2 challenge with no detectable viral titer and viral antigen in the lungs.

**Research in context:** *Evidence before this study:* The spike (S) protein of the SARS-CoV-2 is the major antigen that notably induces neutralizing antibodies to block viral entry. Many COVID-19 vaccines are under development, among them viral vectors expressing the S protein of SARS-CoV-2 exhibit many benefits. Viral vector vaccines have the potential of being used as both live or inactivated vaccines and they can induce Th1 and Th2-based immune responses following different immunization regimens. Additionally, viral vector vaccines can be handled under BSL-2 conditions and they grow to high titers in cell cultures or other species restricted-hosts. For a SARS-CoV-2 vaccine, several viral vectors are being tested, such as adenovirus, measles virus and Modified vaccinia Ankara.

*Added value of this study:* The NDV vector vaccine against SARS-CoV-2 described in this study has advantages similar to those of other viral vector vaccines. But the NDV vector can be amplified in embryonated chicken eggs, which allows for high yields and low costs per dose. Also, the NDV vector is not a human pathogen, therefore the delivery of the foreign antigen would not be compromised by any pre-existing immunity in humans. Finally, NDV has a very good safety record in humans, as it has been used in many oncolytic virus trials. This study provides an important option for a cost-effective SARS-CoV-2 vaccine.

*Implications of all the available evidence:* This study informs of the value of a viral vector vaccine against SARS-CoV-2. Specifically, for this NDV based SARS-CoV-2 vaccine, the existing egg-based influenza virus vaccine manufactures in the U.S. and worldwide would have the capacity to rapidly produce hundreds of millions of doses to mitigate the consequences of the ongoing COVID-19 pandemic.

## Introduction

The unprecedented coronavirus disease 2019 (COVID-19) pandemic caused by severe acute respiratory syndrome coronavirus 2 (SARS-CoV-2) has resulted in ∼16.3 million infections with more than half a million deaths since the end of 2019 as of July 26^th^ 2020, and continues to pose a threat to public health. To mitigate the spread of the virus, social distancing, mask-wearing, and the lockdown of cities, states or even countries were practiced, with a heavy price paid both medically and economically. Unfortunately, due to the relative lack of pre-existing immunity of humans to this virus, no countermeasures will be completely effective without a vaccine. Because of the urgent need for an effective SARS-CoV-2 vaccine, many candidates are being developed using various vaccine platforms, including mRNA vaccines (1, 2), inactivated whole virus vaccines (3), subunit vaccines, DNA vaccines and viral vector vaccines (4). These vaccine candidates are designed to essentially target the spike (S) protein of the SARS-CoV-2 (5), which is the major structural protein displayed on the surface of the SARS-CoV-2. The S protein mediates the entry of the virus via binding to the angiotensin converting enzyme 2 (ACE2) receptor in humans. The S protein is also the most important antigen of the virus that harbors many B cell and T cell epitopes (6-9). Neutralizing antibodies, most of which target the receptor-binding domain (RBD), can be induced by the S protein (9, 10). However, to eventually contain the virus spread worldwide, not only the efficacy, but also the cost and scalability of the vaccine are crucial, especially in low and middle income countries with limited resources.

Here, we report the construction and characterization of Newcastle disease virus (NDV) vectors expressing the SARS-CoV-2 S protein. NDV belongs to the genus of *Avulavirus* in the family of *Paramyxoviridae*, it is an avian pathogen, typically causing no symptoms in humans although mild influenza-like symptoms or conjunctivitis have been described in rare cases. The lentogenic NDV vaccine strain such as the LaSota strain, in addition to be avirulent in birds, has been used as an oncolytic agent and a vaccine vector (11-15). As a large negative strand RNA virus, NDV is stable and well tolerates transgenes into its genome. NDV vectors have been successfully used to express the spike protein of other coronaviruses (16, 17). The NDV platform is also appealing, because the virus grows to high titers in embryonated chicken eggs, which are also used to produce influenza virus vaccines. Humans typically lack pre-existing immunity toward the NDV, which makes the virus preferable over other viral vectors that are human pathogens, such as human adenovirus, measles virus or Modified Vaccinia Ankara (MVA). The lentogenic NDV vector has proven to be safe in humans as it has been tested extensively in human trials (18-20). Most importantly, at low cost, NDV vector vaccines could be generated in embryonated chicken eggs quickly under biosafety level 2 (BSL-2) conditions to meet the vast demand on a global scale. In this study, we have successfully rescued NDV vectors expressing two forms of the spike protein of SARS-CoV-2, the wild type (WT) S and a chimeric version containing the ectodomain (with the polybasic cleavage site deleted) of the spike and the transmembrane domain and cytoplasmic domain of the NDV F (pre-fusion S-F chimera). We have shown that WT S and S-F were well expressed from the NDV as transgenes in infected cells. While both WT S and S-F were displayed on the surface of the NDV particles, the incorporation of the S-F into NDV particles was substantially improved compared to that of the WT S, as expected. A proof of concept study in mice using three live NDV vectors expressing the spike protein (NDV_LS_S, NDV_LS_S-F and NDV_LS/L289A_S-F) showed that high titers of binding and neutralizing antibodies were induced. All three NDV vector vaccines fully protected mice from challenge with a SARS-CoV-2 mouse-adapted strain, showing no detectable viral titers and viral antigens in the lungs at day four post-challenge. To conclude, we have developed promising cost-effective SARS-CoV-2 vaccine candidates using the NDV LaSota strain as the viral vector, which could be generated to high yield under BSL-2 conditions.

## Materials and Methods

### Plasmids

The sequence of the wild type S was amplified from pCAGGS plasmid (21) encoding the codon-optimized nucleotide sequence of the spike gene (GenBank: MN908947.3) of a SARS-CoV-2 isolate by PCR, using primers containing the gene end (GE), gene start (GS) and a Kozak sequences at the 5’ end (22). To construct the S-F chimera, the ectodomain of the S without the polybasic cleavage site (CS, ^682^RRAR^685^ to A) (22) was generated by PCR. A mammalian cell codon-optimized nucleotide sequence of the transmembrane domain (TM) and the cytoplasmic tail (CT) of the NDV LaSota fusion (F) protein was synthesized commercially (gBlock, Integrated DNA technologies). The S ectodomain (no CS) was fused to the TM/CT of F through a GS linker (GGGGS). The sequence was again modified by adding GE, GS and a Kozak sequence at the 5’. Additional nucleotides were added at the 3’ of both inserts to follow the “rule of six”. The transgenes were inserted between the P and M gene of pNDV LaSota (LS) wild type or the L289A (15, 22, 23) mutant (NDV_LS/L289A) antigenomic cDNA by in-Fusion cloning (Clontech). The recombination products were transformed into NEB® Stable Competent E. coli (NEB) to generate NDV_LS_S, NDV_LS_S-F and NDV_LS/L289A_S-F rescue plasmids. The plasmids were purified using QIAprep Spin Miniprep kit (Qiagen) for Sanger sequencing (Macrogen). Maxipreps of rescue plasmids were purified using PureLink™ HiPure Plasmid Maxiprep Kit (Thermo Fisher Scientific).

### Cells

BSRT7 cells stably expressing the T7 polymerase were kindly provided by Dr. Benhur Lee at ISMMS. The cells were maintained in Dulbecco’s Modified Eagle’s medium (DMEM; Gibco) containing 10% (vol/vol) fetal bovine serum (FBS) and 100 unit/ml of penicillin/streptomycin (P/S; Gibco) at 37°C with 5% CO_2_. Vero E6 cells were obtained from American Type Culture Collection (ATCC, CRL-1586). Vero E6 cells were also maintained in DMEM containing 10% FBS with 100 unit/ml P/S at 37 °C with 5% CO_2_.

### Rescue of NDV LaSota expressing the spike protein of SARS-CoV-2

Six-well plates of BSRT7 cells were seeded 3 x 10^5^ cells per well the day before transfection. The next day, a transfection cocktail was prepared consisting of 250 µl of Opti-MEM (Gibco) including 4 µg of pNDV_LS_S or pNDV_LS_S-F or pNDV_LS/L289A_S-F, 2 µg of pTM1-NP, 1 µg of pTM-P, 1 µg of pTM1-L and 2 µg of pCI-T7opt. Thirty µl of TransIT LT1 (Mirus) were added to the plasmid cocktail and gently mixed by pipetting three times and incubated at room temperature (RT) for 30 min. Toward the end of the incubation, the medium was replaced with 1 ml of Opti-MEM. The transfection complex was added dropwise to each well and the plates were incubated at 37°C with 5% CO_2_. Forty-eight hours post transfection, the supernatant and the cells were harvested and briefly homogenized by several strokes with an insulin syringe. Two hundred microliters of the cell/supernatant mixture were injected into the allantoic cavity of 8-to 10-day old specific pathogen free (SPF) embryonated chicken eggs. The eggs were incubated at 37°C for 3 days before being cooled at 4°C overnight. The allantoic fluid was collected and clarified by low-spin centrifugation to remove debris. The presence of the rescued NDV was determined by hemagglutination (HA) assay using 0.5% chicken or turkey red blood cells. The RNA of the positive samples was extracted and treated with DNase I (Thermo Fisher Scientific). Reverse transcriptase-polymerase chain reaction (RT-PCR) was performed to amplify the transgenes. The sequences of the transgenes were confirmed by Sanger Sequencing (Genewiz).

### Immunofluorescence assay (IFA)

Vero E6 cells were seeded onto 96-well tissue culture plates at 2.5 x 10^4^ cells per well. The next day, cells were washed with 100 µl warm phosphate buffered saline (PBS) and infected with 50 µl of allantoic fluid at 37°C for 1h. The inocula were removed and replaced with 100 µl of growth medium. The plates were then incubated at 37°C. Sixteen to eighteen hours after infection, the cells were washed with 100 µl of warm PBS and fixed with 4% methanol-free paraformaldehyde (PFA) (Electron Microscopy Sciences) for 15 min at 4°C. The PFA was discarded, cells were washed with PBS and blocked in PBS containing 0.5% bovine serum albumin (BSA) for 1 hour at 4°C. The blocking buffer was discarded and surface proteins were stained with anti-NDV rabbit serum or SARS-CoV-2 spike receptor-binding domain (RBD) specific human monoclonal antibody CR3022 (24, 25) for 2h at RT. The primary antibodies were discarded, cells were then washed 3 times with PBS and incubated with goat anti-rabbit Alexa Fluor 488 or goat anti-human Alexa Fluor 488 secondary antibodies (Thermo Fisher Scientific) for 1h at RT. The secondary antibodies were discarded, cells were washed again 3 times with PBS and images were captured using an EVOS fl inverted fluorescence microscope (AMG).

### Virus titration

Stocks of NDV expressing the S or S-F proteins were titered using an immunofluorescence assay (IFA). Briefly, Vero cells were seeded onto 96-well (Denville) tissue culture plates at 2.5 x 10^4^ cells/well the day before infection. The next day, five-fold serial dilutions of each virus stocks were prepared in a separate 96-well plate in Opti-MEM (Gibco). Medium in the 96-well plate was removed and the cells were washed with 100 µL of warm PBS. Fifty µL of the virus dilutions were added to each well. The plates were incubated at 37°C for one hour and shaken every 15 minutes to ensure the cells were infected evenly. The inoculum was removed and 100 µL of DMEM containing 10% FBS with 100 unit/ml P/S was added. The plates were incubated at 37°C overnight for 16 to 18 hours. The next day, the media were aspirated off and cells were washed once with 100 µL of warm PBS. IFA was performed as described above to staining NDV surface glycoproteins. Infected fluorescent cells were counted starting from the undiluted wells until a well down the dilution with a countable number of cells was found. The fluorescent cells in the entire well were counted. Titer of the virus (focus forming unit, FFU per ml) was determined by the following formula:

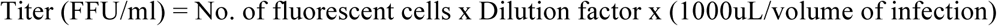

### Virus concentration

Allantoic fluids were clarified by centrifugation at 4,000 rpm using a Sorvall Legend RT Plus Refrigerated Benchtop Centrifuge (Thermo Fisher Scientific) at 4 °C for 30 min. Viruses in the allantoic fluid were pelleted through a 20% sucrose cushion in NTE buffer (100 mM NaCl, 10 mM Tris-HCl, 1 mM ethylenediaminetetraacetic acid (EDTA), pH 7.4) by centrifugation in a Beckman L7-65 ultracentrifuge at 25,000 rpm for 2h at 4°C using a Beckman SW28 rotor (Beckman Coulter, Brea, CA, USA). Supernatants were aspirated and the pellets were re-suspended in PBS (pH 7.4). The protein content was determined using the bicinchoninic acid (BCA) assay (Thermo Fisher Scientific).

### Western Blot

Concentrated virus samples were mixed with Novex™ Tris-Glycine SDS Sample Buffer (2X) (Thermo Fisher Scientific) with NuPAGE™ Sample Reducing Agent (10X) (Thermo Fisher Scientific). The samples were heated at 95 °C for 5 min. Two microgram of concentrated viruses were resolved on 4-20% sodium dodecyl sulfate–polyacrylamide gel electrophoresis (SDS-PAGE) gels (Biorad) using the Novex™ Sharp Pre-stained Protein Standard (Thermo Fisher Scientific) as the marker. Proteins were transferred onto polyvinylidene difluoride (PVDF) membrane (GE healthcare). The membrane was blocked with 5% dry milk in PBS containing 0.1% v/v Tween 20 (PBST) for 1h at RT. The membrane was washed with PBST on a shaker 3 times (10 min at RT each time) and incubated with primary antibodies diluted in PBST containing 1% BSA overnight at 4°C. To detect the spike protein of SARS-CoV-2, a mouse monoclonal antibody 2B3E5 kindly provided by Dr. Thomas Moran at ISMMS was used, while the HN protein was detected by a mouse monoclonal antibody 8H2 (MCA2822, Bio-Rad). The membranes were then washed with PBST on a shaker 3 times (10 min at RT each time) and incubated with sheep anti-mouse IgG linked with horseradish peroxidase (HRP) diluted (1:2,000) in PBST containing 5% dry milk for 1h at RT. The secondary antibody was discarded and the membranes were washed with PBST on a shaker 3 times (10 min at RT each time). Pierce™ ECL Western Blotting Substrate (Thermo Fisher Scientific) was added to the membrane, the blots were imaged using the Bio-Rad Universal Hood Ii Molecular imager (Bio-Rad) and processed by Image Lab Software (Bio-Rad).

### Mice immunizations

Ten-week old female BALB/cJ mice (Jackson Laboratories) were used. Experiments were performed in accordance with protocols approved by the Icahn School of Medicine at Mount Sinai Institutional Animal Care and Use Committee (IACUC). Mice were divided into 9 groups (n=5) receiving four different concentrated live viruses at two doses (10 µg and 50 µg) intramuscularly (i.m). Specifically, group 1 (10 µg per mouse) and 2 (50 µg per mouse) were given wild type NDV_LS; group 3 (10 µg per mouse) and 4 (50 µg per mouse) received NDV_LS_S; group 5 (10 µg) and 6 (50 µg) received NDV_LS_S-F and group 7 (10 µg per mouse) and 8 (50 µg per mouse) received NDV_LS/L289A_S-F. Group 9 given PBS was used as the negative controls. A prime-boost immunization regimen was used for all the groups in a 3-week interval.

### Enzyme linked immunosorbent assay (ELISA)

Immunized mice were bled pre-boost and 8 days after the boost. Sera were isolated by low-speed centrifugation. To perform ELISAs, Immulon 4 HBX 96-well ELISA plates (Thermo Fisher Scientific) were coated with 2 µg/ml of recombinant trimeric S protein (50 µL per well) in coating buffer (SeraCare Life Sciences Inc.) overnight at 4°C (21). The next day, all plates were washed 3 times with 220 µL PBS containing 0.1% (v/v) Tween-20 (PBST) and 220 µL blocking solution (3% goat serum, 0.5% dry milk, 96.5% PBST) was added to each well and incubated for 1h at RT. Mouse sera were 3-fold serially diluted in blocking solution starting at 1:30 followed by a 2 h incubation at RT. ELISA plates were washed 3 times with PBST and 50 µL of sheep anti-mouse IgG-horseradish peroxidase (HRP) conjugated antibody (GE Healthcare) was added at a dilution of 1:3,000 in blocking solution. Then, plates were again incubated for one hour at RT. Plates were washed 3 times with PBST and 100 µL of o-phenylenediamine dihydrochloride (SigmaFast OPD, Sigma) substrate was added per well. After 10 min, 50 µL of 3M hydrochloric acid (HCl) was added to each well to stop the reaction and the optical density (OD) was measured at 492 nm on a Synergy 4 plate reader (BioTek). An average of OD values for blank wells plus three standard deviations was used to set a cutoff for plate blank outliers. A cutoff value was established for each plate that was used for calculating the endpoint titers. The endpoint titers of serum IgG responses was graphed using GraphPad Prism 7.0.

### SARS-CoV-2 challenge in mice

The SARS-CoV-2 challenge was performed at the University of North Carolina by Dr. Ralph Baric’s group in a Biosafety Level 3 (BSL-3) facility. Mice were challenged 11 days after the boost using a mouse adapted SARS-CoV-2 strain at 10^4^ plaque forming unit (PFU) intranasally (i.n) under ketamine/xylazine anesthesia as described previously (1, 26).

### Lung titers

Lung lobes of mice were collected and homogenized in PBS. A plaque assay was performed to measure viral titer in the lung homogenates as described previously (1, 26). Geometric mean titers of plaque forming units (PFU) per lobe were calculated using GraphPad Prism 7.0.

### Micro-neutralization assay

All neutralization assays were performed in the biosafety level 3 (BSL-3) facility following institutional guidelines as described previously (21, 27). Briefly, serum samples were heat-inactivated at 56°C for 60 minutes prior to use. 2X minimal essential medium (MEM) supplemented with glutamine, sodium biocarbonate, 4- (2- hydroxyethyl)1- piperazineethanesulfonic acid (HEPES), and antibiotics P/S was used for the assay. Vero E6 cells were maintained in culture using DMEM supplemented with 10% fetal bovine serum (FBS). Twenty-thousands cells per well were seeded the night before in a 96-well cell culture plate. 1X MEM was prepared from 2X MEM and supplemented with 2% FBS. Three-fold serial dilutions starting at 1:20 of pooled sera were prepared in a 96-well cell culture plate and each dilution was mixed with 600 times the 50% tissue culture infectious dose (TCID_50_) of SARS-CoV-2 (USA-WA1/2020, BEI Resources NR-52281). Serum-virus mixture was incubated for 1h at room temperature. Virus-serum mixture was added to the cells for 1h and kept in a 37°C incubator. Next, the virus-serum mixture was removed and the corresponding serum dilution was added to the cells with addition 1X MEM. The cells were incubated for 2 days and fixed with 100 µL 10% formaldehyde per well for 24 h before taken out of the BSL-3 facility. The staining of the cells was performed in a biosafety cabinet (BSL-2). The formaldehyde was carefully removed from the cells. Cells were washed with 200 µL PBS once before being permeabilized with PBS containing 0.1% Triton X-100 for 15 min at RT. Cells were washed with PBS and blocked in PBS containing 3% dry milk for 1h at RT. Cells were then stained with 100 µL per well of a mouse monoclonal anti-NP antibody (1C7), kindly provided by Dr. Thomas Moran at ISMMS, at 1µg/ml for 1h at RT. Cells were washed with PBS and incubated with 100 µL per well Anti-mouse IgG HRP (Rockland) secondary antibody at 1:3,000 dilution in PBS containing 1% dry milk for 1h at RT. Finally, cells were washed twice with PBS and the plates were developed using 100 µL of SigmaFast OPD substrate. Ten minutes later, the reactions were stopped using 50 µL per well of 3M HCI. The OD 492 nM was measured on a Biotek SynergyH1 Microplate Reader. Non-linear regression curve fit analysis (The top and bottom constraints are set at 100% and 0%) over the dilution curve was performed to calculate 50% of inhibitory dilution (ID_50_) of the serum using GraphPad Prism 7.0.

### Immunohistochemistry (IHC)

The lung lobes of mice were perfused and fixed in 10% phosphate buffered formalin for 7 days before transferred out of the BSL-3 facility. The fixed lungs were paraffin embedded, and sectioned at 5µm for immunohistochemistry (IHC) staining (HistoWiz). IHC was performed using a rabbit SARS-CoV-2 nucleocapsid (N) protein (NB100-56576, Novus Biologicals). Slides were counter stained with hematoxylin. All slides were examined by a board-certified veterinary pathologist (HistoWiz).

## Results

### Design and rescue of NDV LaSota expressing the spike protein of SARS-CoV-2

For protective immunity, the S protein is the most important antigen of SARS-CoV-2. To express S antigen by the NDV LaSota vaccine strain, we designed two constructs. One is the wild type spike (S), the other is the spike-F chimera (S-F). The S-F consists of the ectodomain of the S, in which the polybasic cleavage site ^682^RRAR^685^ is removed by deleting the three arginines to stabilize the protein in its pre-fusion conformation (21). Importantly, to increase membrane-anchoring of the spike on the surface of the NDV virions, we replaced the transmembrane domain (TM) and cytoplasmic tail (CT) of the spike with those from the fusion (F) protein of NDV (Fig. 1A)(28). The nucleotide sequences of each construct were inserted between the P and M genes of the antigenomic cDNA of WT NDV LaSota strain and/or NDV LaSota/L289A mutant strain, in which the mutation L289A in the F protein supports HN independent fusion (23). The latter NDV mutant has been safely used in humans (15) (Fig. 1B). NDV expressing the spike proteins were rescued by transient transfection of BSRT7 cells followed by amplification in embryonated chicken eggs. All the viruses expressing the S or S-F grew to high titers (∼10^8^ FFU/ml) in embryonated chicken eggs (Fig. 1C), which is advantageous for the development of a low-cost vaccine.

**Figure 1.**
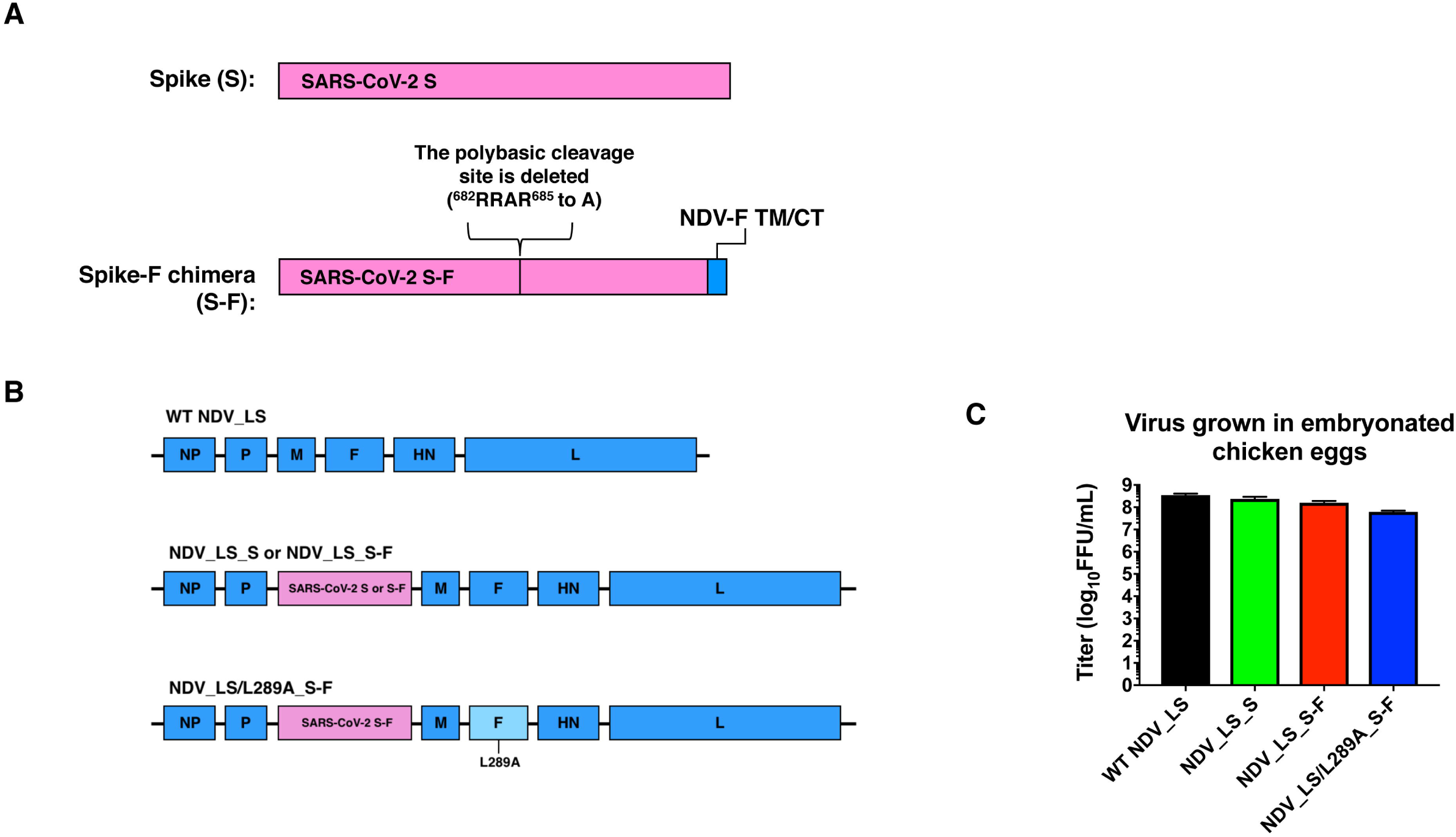
NDV vectors expressing the spike protein of SARS-CoV-2. (A) Two forms of spike proteins expressed by NDV. Spike (S) has the wild type amino acid sequence. The Spike-F chimera (S-F) consists of the ectodomain of S without the polybasic cleavage site and the transmembrane domain (TM) and cytoplasmic tail (CT) of the F protein from NDV. (B) Illustration of genome structures of wild type NDV LaSota (WT NDV_LS), NDV expressing the S or S-F in the wild type LaSota backbone (NDV_LS_S or NDV_LS_S-F) or NDV expressing the S-F in the L289A mutant backbone (NDV_LS/L289A_S-F). The L289A mutation supports the HN-independent fusion of the F protein. (C) Titers of NDV vectors grown in embryonated chicken eggs. The rescued viruses were grown in 10-day old embryonated chicken eggs for 2 or 3 days at 37 °C at limiting dilutions. The peak titers of each virus were determined by immunofluorescence assay (IFA).

### The spike protein is incorporated into NDV particles

To validate the expression of S and S-F as transgenes, Vero E6 cells were infected with WT NDV or NDV expressing the S or S-F. The surface of the cells was stained with anti-NDV rabbit serum or spike-specific monoclonal antibody CR3022 that recognizes the RBD. We confirmed that only NDV expressing the S or S-F showed robust expression of the spike on the cell surface, while NDV proteins were detected in all virus-infected cells (Fig. 2A). This demonstrates that S and S-F are successfully expressed by the NDV. To examine the incorporation of the S and S-F into the NDV virions, we concentrated the NDV_LS_S, NDV_LS_S-F and NDV_LS/L289A_S-F through a 20% sucrose cushion. The pellets were re-suspended in PBS. The WT NDV_LS was prepared the same way and was used as the negative control. The protein content of each concentrated virus was determined by BCA assay. Two micrograms of each virus was resolved on an SDS-PAGE. A Western blot was performed to examine the abundance of the spike using mouse monoclonal antibody 2B3E5 that binds to a linear epitope of the S1 protein. The expression of the NDV viral hemagglutinin-neuraminidase (HN) protein was also shown as an internal control of the concentrated viruses (Fig. 2B). As expected, both S and S-F incorporated into the NDV particles. Of note, the WT S harboring the polybasic cleavage site (CS) was completely cleaved showing only the S1, while the S-F was maintained at its pre-fusion S0 stage. Importantly, the S-F expressed either by the WT or L289A NDV_LS backbone exhibited superior incorporation into the virions over the WT S shown by much higher abundance of S-F than S1 cleaved from the WT S (Fig. 2B). This confirms that the TM/CT of F in the S-F chimera indeed facilitates the membrane-anchoring of the spike as expected. Since the anti-NDV rabbit sera completely neutralize focus formation of these three NDV vectors vaccines, and for the fact, that the S-F constructs don’t have a polybasic cleavage site, it is unlikely the expression of the transgenes alters the tropism of these viruses.

**Figure 2.**
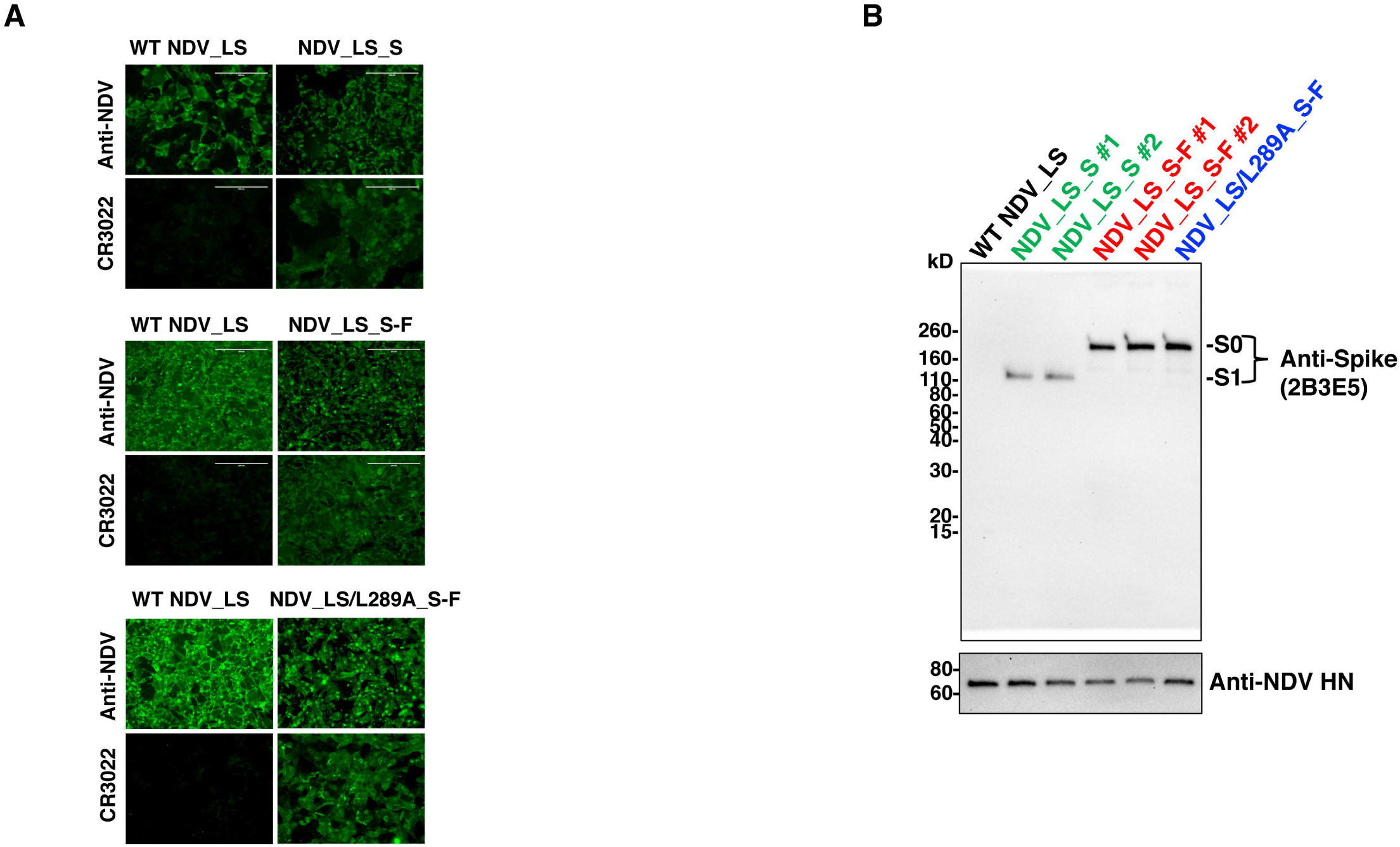
Expression of spike protein in infected cells and NDV particles. (A) Expression of the S and S-F protein in infected cells. Vero E6 cells were infected with three NDV vectors encoding the S or S-F for 16 to 18 hours. A WT NDV control was included. The next day, cells were fixed with methanol-free paraformaldehyde. Surface proteins were stained with anti-NDV rabbit serum or a spike receptor-binding domain (RBD)-specific monoclonal antibody CR3022. (B) Incorporation of S and S-F into NDV particles. Three NDV vectors expressing the S or S-F including the NDV_LS_S (green), NDV_LS_S-F (red) and NDV_LS/L289A_S-F (blue) were concentrated through a 20% sucrose cushion. Two clones were shown for NDV_LS_S and NDV_LS_S-F. The concentrated WT NDV expressing no transgenes was used as a control. Two micrograms of each concentrated virus were resolved on a 4-20% SDS-PAGE, the spike protein and NDV HN protein were detected by western blot using an anti-spike 2B3E5 mouse monoclonal antibody and an anti-HN 8H2 mouse monoclonal antibody.

### Immunization of mice with NDV LaSota expressing the spike protein elicited potently binding and neutralizing antibodies

To evaluate the immunogenicity of our NDV vectors expressing the S or S-F as vaccine candidates against SARS-CoV-2, a proof of principle study was performed in mice. Specifically, BALB/c mice were immunized with live NDV_LS_S, NDV_LS_S-F and NDV_LS/L289A_S-F intramuscularly, as live NDV barely replicates in the muscle and causes no symptoms in mammals. Here, we used a prime-boost immunization regimen in a three-week interval. Mice were bled pre-boost (after prime) and 8 days after the boost for *in vitro* serological assays (Fig. 3A). Two doses (10 µg and 50 µg) of each NDV construct including NDV_LS_S (group 3 and 4), NDV_LS_S-F (group 5 and 6) and NDV_LS_S-F (group 7 and 8) were tested as shown in Fig. 3A. Animals vaccinated with WT NDV expressing no transgenes (group 1 and 2) were used as vector-only controls. Mice receiving only the PBS (group 9) were used as negative controls. Mouse sera from the two bleedings were harvested. Serum IgG titers and neutralizing antibody titers were measured by ELISAs and microneutralization assays, respectively. To perform ELISA, full-length trimeric spike protein was coated onto ELISA plates. The endpoint titers of serum IgG were used as the readout (Fig. 3B). After one immunization, all the NDV constructs expressing the spike protein elicited S-binding antibodies, whereas WT NDV constructs and PBS controls show negligible antibody binding signals. The second immunization significantly increased the antibody titers around 1 week after the boost without showing significant difference among the three NDV constructs (Fig. 3B). The neutralizing activity of the antibodies was measured in a microneutralization assay using the USA-WA1/2020 SARS-CoV-2 strain. Pooled sera from each group were tested in a technical duplicate. The ID_50_ value was calculated as the readout of neutralizing activity of post-boost sera (Day 29). Sera from all vaccinated groups showed neutralizing activity. The neutralization titer of sera from the NDV_LS_S high-dose (50 µg) vaccination group (ID_50_ ≈ 444) appeared to be slightly higher than that from the low-dose (10 µg) vaccination group (ID_50_ ≈178). No substantial difference was observed between the low-dose and high-dose groups using the NDV_LS_S-F and NDV_LS/L289A_S-F constructs (Fig. 3C), the neutralization titers of which are comparable to that of the NDV_LS_S high-dose (50 µg) group. To summarize, all the NDV vectors that were engineered to express the S or S-F elicited high titers of binding and neutralizing antibodies in mice. The WT S and S-F constructs appeared to exhibit similar immunogenicity when expressed by live NDV vectors that were given intramuscularly to mice.

**Figure 3.**
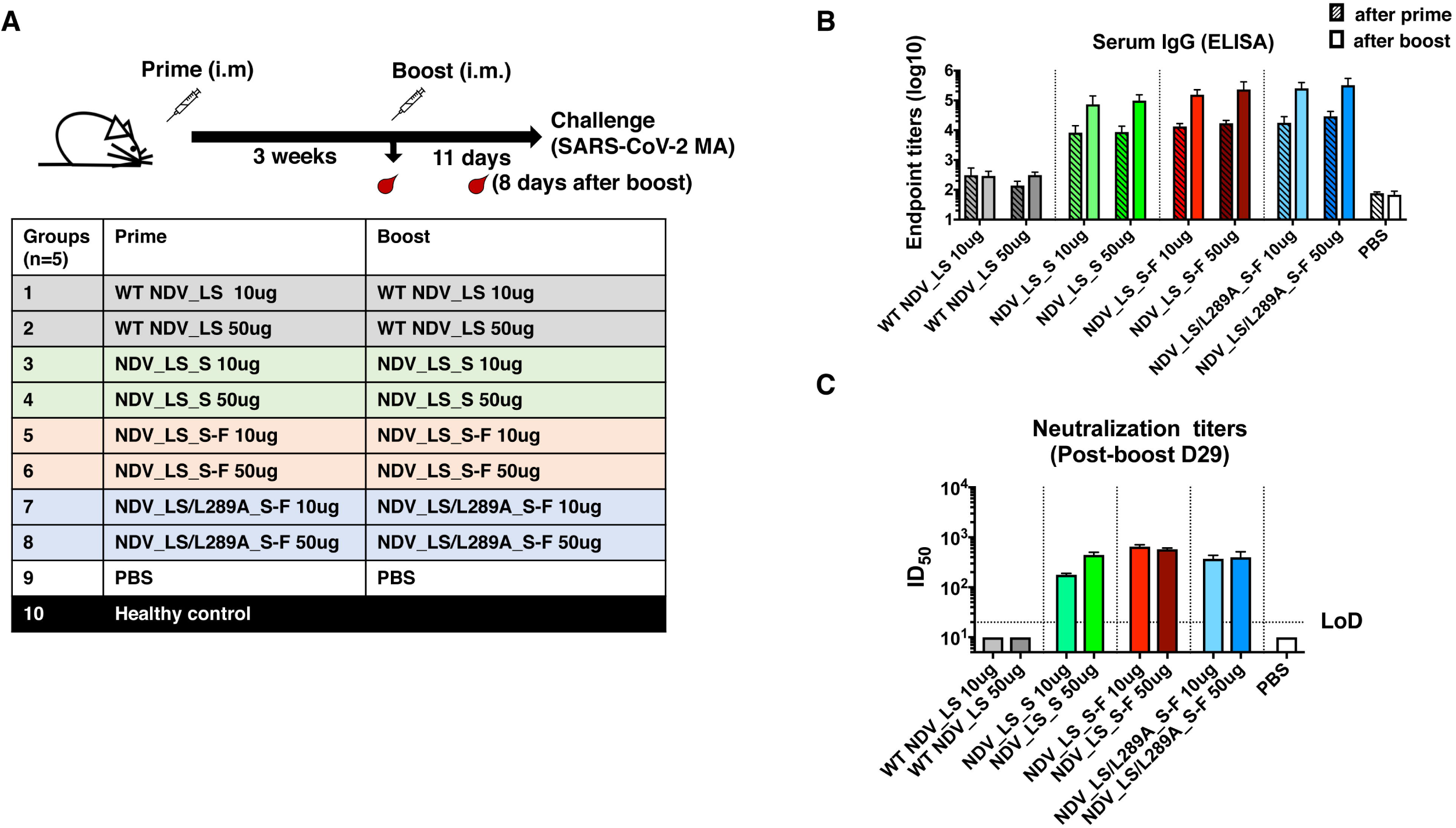
NDV vector vaccines elicit high titers of binding and neutralizing antibodies in mice. (A) Vaccination groups and regimen. A prime-boost vaccination regimen was used with a three-week interval. Mice were bled pre-boost and 8 days after the boost. Mice were challenged with a mouse-adapted SARS-CoV-2 MA strain 11 days after the boost. A total of ten groups of mice were used in a vaccination and challenge study. Group 1 (10 µg) and 2 (50 µg) received the WT NDV; Group 3 (10 µg) and 4 (50 µg) received the NDV_LS_S; Group 5 (10 µg) and 6 (50 µg) received NDV_LS_S-F; Group 7 (10 µg) and 8 (50 µg) received NDV_LS/L289A_S-F; Group 9 received PBS as negative controls. An age-matched healthy control group 10 was provided upon challenge. (B) Spike-specific serum IgG titers measured by ELISAs. Sera from animals at 3 weeks after-prime (patterned bars) and 8 days after-boost (solid bars) were isolated. Serum IgG was measured against a recombinant trimeric spike protein by ELISAs. The endpoint titers were calculated as the readout for ELISAs. (C) Neutralization titers of serum antibodies. Sera from 3 weeks after-prime and 8 days after-boost were pooled within each group. Technical duplicates were performed to measure neutralization activities of serum antibodies using a USA-WA1/2020 SARS-CoV-2 strain. The ID50 value was calculated as the readout of the neutralization assay. For the samples (WT NDV and PBS groups) showing no neutralizing activity in the assay, an ID50 of 10 was given as the starting dilution of the sera is 1:20 (LoD: limit of detection).

### Immunization with NDV LaSota expressing the spike proteins protects mice from challenge with a mouse-adapted SARS-CoV-2

To assess *in vivo* activity of S-specific antibodies induced by the NDV constructs as well as potential cell-mediated protection, we took advantage of a mouse – adapted SARS-CoV-2 strain that replicates efficiently in BALB/c mice (1, 26). The immunized mice were challenged with 10^4^ PFU of the mouse-adapted SARS-CoV-2 at day 11 after the boost, and viral titers in the lungs at day 4 post –challenge were measured. Mice receiving WT NDV and PBS exhibited high viral titers in the lung, while all the groups given NDV expressing the S or S-F showed no detectible viral load in the lung (Fig. 4A). The lungs of infected mice were fixed in 10% neutral buffered formalin for IHC staining using an anti-SARS-CoV-2 NP antibody. The IHC staining showed that the SARS-CoV-2 NP protein was largely detected in the lungs of mice that received NDV_LS WT or PBS. The SARS-CoV-2 NP was absent in the lungs of mice vaccinated with the three NDV constructs expressing the S or S-F protein (Fig. 4B). These data demonstrated that the all three NDV vector vaccines could efficiently prevent SARS-CoV-2 infection in a mouse model.

**Figure 4.**
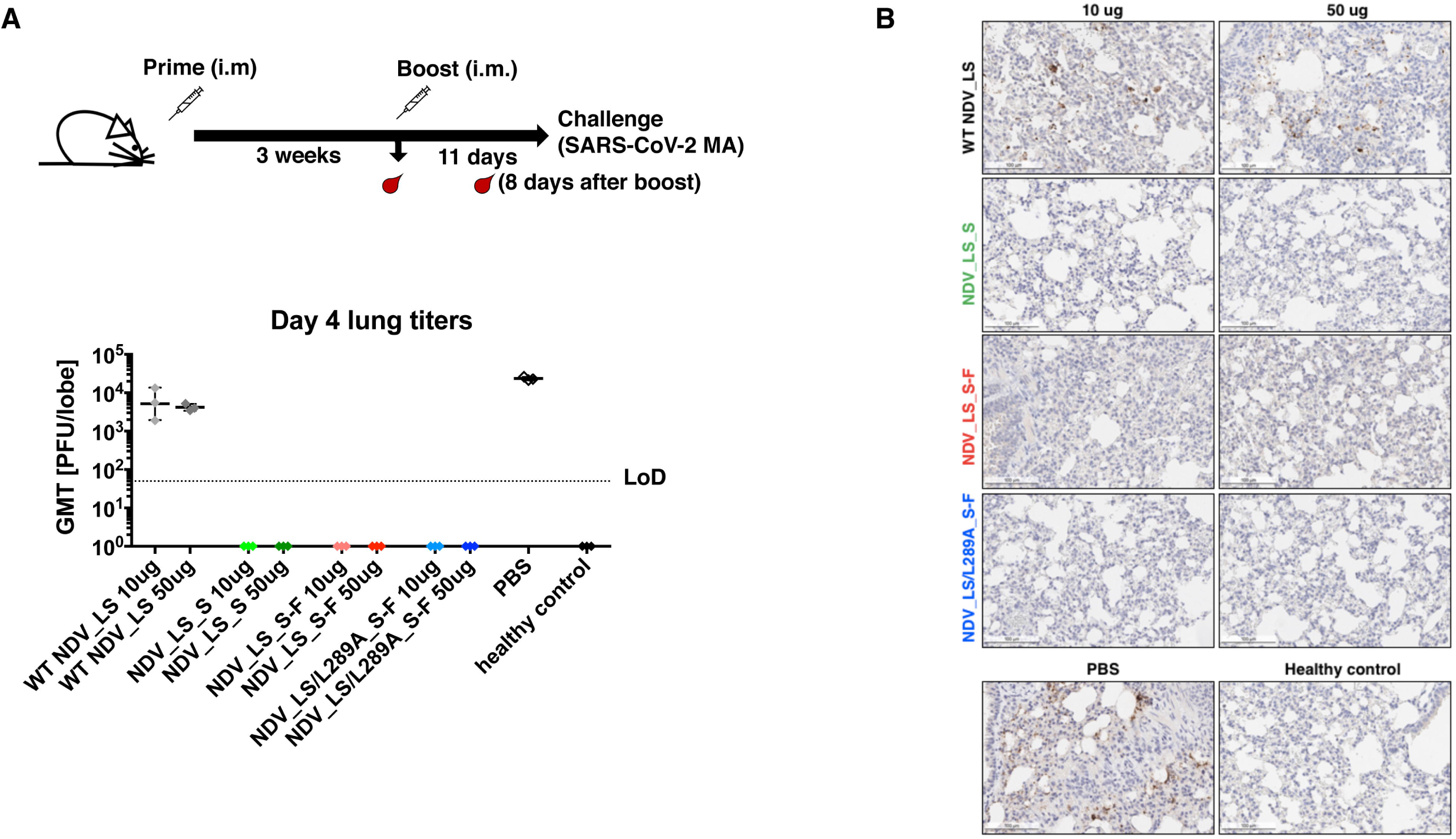
NDV vector vaccines protected mice from the SARS-CoV-2 challenge. (A) Viral titers in the lungs. All mice were infected intranasally with 10^4^ PFU SARS-CoV-2 MA strain except the healthy control group, which was mock infected with PBS. At day 4 post-challenge, lungs were collected and homogenized in PBS. Viral titers in the lung homogenates were determined by plaque assay. Plaque-forming units (PFU) per lung lobe was calculated. Geometric mean titer was shown for all the groups. LoD: limit of detection. (B) Immunohistochemistry (IHC) staining of lungs. A SARS-CoV-2 NP specific antibody was used for IHC to detect viral antigens. Slides were counterstained with hematoxylin. A presentative image was shown for each group. The brown staining indicates the presence of NP protein of SARS-CoV-2.

## Discussion

The consequences of the ongoing COVID-19 pandemic since the end of 2019 are disheartening. With the high transmissibility of the culprit, SARS-CoV-2, and the lack of substantial pre-existing immunity of humans to this virus, many people have succumbed to COVID-19, especially the elderly and people with underlying health conditions. With both therapeutic and prophylactic countermeasures (29-31) still under rapid development, no currently available treatment appears to be effective enough for an over-burdened health care system with limited resources. A vaccine is needed to prevent or at least attenuate the symptoms of COVID-19. As many vaccine candidates are being tested in pre-clinical or clinical studies, a vaccine for cost effective production in low-and middle-income countries has not yet been developed and is still very much in need. Also, the vaccination of small numbers of high-income populations who can afford the vaccine would not efficiently prevent the spreading of the disease in the global population. In this report, we describe promising viral vector vaccine candidates based on NDV expressing the major antigen of SARS-CoV-2. The NDV vectors were engineered to express either the wild type S or a prezfusion spike with improved membrane anchoring (S-F). These NDV vector vaccines showed robust growth in embryonated chicken eggs despite the fact that a large transgene is inserted into the NDV genome. Importantly, the spike protein is successfully expressed in infected cells, and the S-F construct exhibited superior incorporation into NDV particles, which could potentially be used as an inactivated virus vaccine as well.

In a proof of principle study, mice receiving live NDV vector vaccines twice intramuscularly have developed high levels of spike-specific antibodies that are neutralizing. Mice given the NDV vector expressing S or S-F were protected equally well against the challenge of a mouse-adapted SARS-CoV-2 strain showing no detectable infectious virus or viral antigens in the lungs, while high viral titers were observed in the lungs of mice given the WT NDV expressing no transgenes or PBS. In this study, we did not see significant dose-dependent antibody responses, which was similar to what was observed for different doses (100 µg and 250 µg) of mRNA vaccine in a human trial (2). It could be that we did not measure the peak antibody responses due to the problem of having to transfer mice to the University of North Carolina for the challenge study, or an antibody response ceiling was reached with the low dose of 10 µg concentrated virus in mice. In the present study, cellular immunity was not measured, however, this will be of interest to investigate in the future studies. Nevertheless, this study strongly supports that the NDV vector vaccines are promising, as they are expressing immunogenic spike proteins of SARS-CoV-2 inducing high levels of protective antibodies. Unlike other viral vectors that humans might be exposed to, the NDV vector would deliver the spike antigen more efficiently without encountering pre-existing immune responses in humans. Importantly, NDV vector vaccines are not only cost-effective with respect to large scale manufacturing but can also be produced under BSL-2 conditions using influenza virus vaccine production technology. In summary, NDV vector SARS-CoV-2 vaccines are a safe and immunogenic alternative to other SARS-CoV-2 vaccines that can be produced using existing infrastructure in a cost-effective way.

## Contributors

Conceptualization and design, P.P.; methodology, P.P., W.S., S.R.L, S.M., Y.L, S.S., J.O, F.A., A.G.S, F.K., R.S.B; investigation and data analysis, P.P., W.S., S.R.L, S.M., Y.L, S.S., J.O, F.A., A.S, K.H.D, A.G.S, F.K., R.S.B; first draft of manuscript, P.P. and W.S.; manuscript review and editing, all authors; funding, P.P. and anonymous philanthropist.

## Acknowledgement

We thank Dr. Benhur Lee to kindly share the BSRT7 cells. We also thank Dr. Thomas Moran for the 2B3E5 and 1C7 antibodies. This work was partially supported by an NIAID funded Center of Excellence for Influenza Research and Surveillance (CEIRS, HHSN272201400008C, P.P.) and a grant from an anonymous philanthropist to Mount Sinai (PP). Work in the Krammer laboratory was also partially supported by the NIAID Centers of Excellence for Influenza Research and Surveillance (CEIRS) contract HHSN272201400008C (FK) and by the Collaborative Influenza Vaccine Innovation Centers (CIVIC) contract 75N93019C00051 (FK), the generous support of the JPB foundation, the Open Philanthropy Project (#2020-215611) and other philanthropic donations. RSB was supported by NIH U01 AI149644.

